# Extinct species identification from Upper Pleistocene bone fragments not identifiable from their osteomorphological studies by proteomics analysis

**DOI:** 10.1101/2020.10.06.328021

**Authors:** Fabrice Bray, Stéphanie Flament, Grégory Abrams, Dominique Bonjean, Kévin Di Modica, Christian Rolando, Caroline Tokarski, Patrick Auguste

## Abstract

The ancient preserved molecules offer the opportunity to gain a better knowledge on the biological past. In recent years, bones proteomics has become an attractive method to study the animal biological origin, extinct species and species evolution as an alternative to DNA analysis which is limited by DNA amplification present in ancient samples and its contamination. However, the development of a proteomic workflow remains a challenge. The analysis of fossils must consume a low quantity of material to avoid damaging the samples. Another difficulty is the absence of genomic data for most of the extinct species. In this study, a proteomic methodology was applied to mammalian bones of 130,000 years old from the earlier Upper Pleistocene site of Scladina Cave (Belgium). Starting from 5 milligram samples, our results show a large majority of detected peptides matching collagen I alpha 1 and alpha 2 proteins with a sequence coverage up to 60%. Using sequence homology with modern sequences, a biological classification was successfully achieved and the associated taxonomic ranks to each bone were identified consistently with the information gained from osteomorphological studies and palaeoenvironmental and palaeodietary data. Among the taxa identified are the Felidae family, Bovinae subfamily, Elephantidae family and the Ursus genus. Amino acid substitutions on the collagens were identified providing new information on extinct species sequences and also helping in taxonomy-based clustering. Considering samples with no osteomorphological information, such as two bone retouchers, proteomics successfully identified the bovidae and ursidae families providing new information to the paleontologists on these objects. Combining osteomorphology studies and amino acid variations identified by proteomics, one retoucher was identified to be potentially from the *Ursus spelaeus* species.

## 1. Introduction

The study of bones by mass spectrometry (MS) has become an increasingly used method. Indeed, bones are real safes protecting their constitutive proteins. MS proved to be a robust and accurate method allowing the identification of proteins, their biological origins and their modifications (Cleland and Schroeter, 2018, Dallongeville, et al., 2016, Vinciguerra, et al., 2016, Welker, 2018) overpassing DNA analysis considering the longer temporal scales (Cappellini, et al., 2018, Demarchi, et al., 2016). For example, the identification of collagen I from *Brachylophosaurus canadensis* dinosaurs has been confirmed (Schroeter, et al., 2017) and collagen sequences from fossil *camels*, *Camelops* and c.f. *Paracamelus*, from the Arctic and sub-Arctic of Plio-Pleistocene North America has been reported (Buckley, et al., 2019). Another examples are the palaeoproteomics studies of *Gigantopithecus blacki* dated between the Early Pleistocene and the late Middle Pleistocene (Welker, et al., 2019) and Eurasian Rhinocerotidae of the Pleistocene epoch (Cappellini, et al., 2019).

Type I collagen is the major protein of bones. It is constituted by the association of two alpha 1 and one alpha 2 chains arranged in a super helix. The sequence of this protein is characterized by regular occurrences of glycine, proline and hydroxyproline (Shoulders and Raines, 2009). Non-collagenous proteins (NCPs) represent a smaller part of bone proteins (Young, et al., 1992). Type I collagen is the main protein identified by mass spectrometry in ancient bones.

Palaeoproteomic analysis of bones consists of a first protein/peptide preparation step, generally starting from several tens to hundreds milligrams of bones, and their analysis and identification by mass spectrometry and dedicated bioinformatics tools. The starting material amount is low in comparison to other techniques usually applied to bone analysis such as ^14^C dating (from several hundreds of milligrams to gram). The first step of a classical palaeoproteomic analysis is the demineralization which is performed most often by an acidic solution and seldom by a basic solution (Schroeter, et al., 2016). This step is well-established for decalcification and solubilization. The extracted proteins are then digested into peptides by an enzyme, most often the trypsin. Several digestion methods have been reported such as liquid digestion (Horn, et al., 2019, Sawafuji, et al., 2017), filter assisted sample preparation (FASP) (Cappellini, et al., 2014, Kostyukevich, et al., 2018) and solid digestion of demineralized bones (Cleland, 2018b). Recently was proposed a single-pot solid-phase-enhanced sample preparation (SP3) (Cleland, 2018a) which is an effective method to enable collagen and NCPs analyses along with the removal of coextracted humic compounds. Another method proposes to study the adhering collagen from the storage plastic bags (McGrath, et al., 2019). This technique allows studying the object without affecting the integrity of the artifact. Multi-protease approaches were also described to increase the protein sequence coverage (Lanigan, et al., 2020). Another recent studies in paleoproteomics also propose a digestion-free approach to characterize ancient remains (Cappellini, et al., 2019, Welker, et al., 2019).

Two techniques are used to identify the peptides extracted from bones: ZooMS and shotgun analysis based on LC separation hyphenated with ESI-MS. Developed in 2009, ZooMS (Buckley, et al., 2009) is based on peptide mass fingerprint analysis using MALDI-TOF MS. It allows taxonomic discrimination between a wide range of mammalian bones by the identification of peptides variants between species. ZooMS method has resulted in the identification of various species from ancient material such as hominin (Brown, et al., 2016), marine mammals (Hofman, et al., 2018), other mammals such as sheep, goat (Birch, et al., 2019, Brandt, et al., 2018), kangaroos (Buckley, et al., 2017a) from bones but also from derived material such as bones tools (Desmond, et al., 2018, Martisius, et al., 2020, McGrath, et al., 2019). Considering shotgun analysis, the amino acid sequences of the peptides are identified allowing thus an accurate identification of proteins including the species-related amino acids variants. The first sequencing experiments were proposed in the early 2000s on bone osteocalcin (Nielsen-Marsh, et al., 2002, Nielsen-Marsh, et al., 2005, Ostrom, et al., 2006, Ostrom, et al., 2000). Early 2010s showed the first application of shotgun analysis on a 43,000 year old woolly mammoth bone (Cappellini, et al., 2011) resulting in the identification of a hundred proteins. Since, the method has been extensively applied to the study of other extinct species such as *Castoroides ohioensis* (Cleland, et al., 2016), *Bison latifrons* (Hill, et al., 2015), ancient birds (Horn, et al., 2019), dinosaurs (Schroeter, et al., 2017). Along with the study of extinct species, phylogeny and evolutionary history are studied. Palaeoproteomics applied to the study of endemic South American ‘ungulates’ (Buckley, 2015), Darwin’s South American ungulates (Welker, et al., 2015), ancient hominin specimens (Welker, et al., 2016), archaeological marine turtles (Harvey, et al., 2019) and tree sloth (Presslee, et al., 2019) are several of these applications.

We present here a robust proteomics workflow to study extinct species from Scladina Cave (Andenne, Belgium) dated from earlier Upper Pleistocene (130,000 years B.C.). The method was applied to 5 milligrams of starting material. Both ancient bones and retouchers were studied. Our analyses resulted in a successful characterization of biological classification and associated taxonomic ranks to each bone consistently with the information gained from morphological studies. Taxa identifications were also provided for the retouchers (not identified from their morphology) in agreement with the fauna found on the site. It can be pointed out that during this study, amino acid substitutions on the collagens were identified providing new information on extinct species sequences and also helping in taxonomy-based clustering.

## 2. Materials and methods

### 2.1 Samples

Ten bones from the archaeological excavation of Scladina Cave were selected from different geological layers (Units 4B, 5, 6A, 6B, 6C, 4B). Scladina Cave is located 400 m southwest of the village of Sclayn located in Belgium between Andenne and Namur, close to the Meuse River. Several dating methods were established to estimate the site to be 130,000 years old (Otte, et al., 1983, Pirson, et al., 2008, Pirson, et al., 2014). The species of 8 bones on 10 were identified through their anatomy. The 2 remaining ones used as retouchers or “lissoirs” (smoothers) were from unknown or hypothetic origins. These two last bones are of particular interest because they are directly associated with retouchers by refitting. Bone. The Table 1 is referencing the 10 studied samples with archaeological information. Photographs of the studied bones are presented in Supplementary data file 1 Fig. S1.1 to S1.10.

**Table 1.** Bones samples from the Scladina Cave (Sclayn, Belgium).

| Excavation year | Excavation number | Geological Unit | Location (square) | Description | Species | Sample name |
| --- | --- | --- | --- | --- | --- | --- |
| 1983 | 236-52 | 5 | E13 | Calcaneus | *Panthera spelaea* | Sc-1 |
| 2011 | 362-7 | 6C-LAGJ | G16 | Metatarsal II | *Panthera spelaea* | Sc-2 |
| 1983 | 407-93 | 5 | E12 | Metatarsal III | *Panthera spelaea* | Sc-3 |
| 1981 | 127-3 | 6A | C4 | Humerus | *Panthera pardus* | Sc-4 |
| 2013 | 46-2 | 6B | C37 | Metatarsal III | *Ursus spelaeus* | Sc-5 |
| 2013 | 44-8 | 6A | D37 | Metatarsal I | *Ursus thibetanus* | Sc-6 |
| 2003 | 590-2 | 6A | B36 | Metatarsal III | *Ursus thibetanus* | Sc-7 |
| 1995 | 430-97 | 4B | C36 | Rib | *Mammuthus* sp | Sc-8 |
| 1987 | 52 | 5 | I22 | Retoucher | Unknown | Sc-9 |
| 1985 | F16-17 | 5 | F16 | Retoucher | Unknown | Sc-10 |

### 2.2 Chemicals and biochemicals

All aqueous solutions were prepared from ultrapure grade water obtained by water filtration with a two stages Millipore system (Milli-Q® Academic with a cartouches Q-Gard 1 and Progard 2, Merck Millipore, Burlington, Massachusetts, United States). All chemicals, biochemicals and solvents were purchased from Sigma-Aldrich (Saint-Louis, Missouri, USA) and used without purification. All solvents were MS analytical grade.

### 2.3 Protein extraction from bones and delipidation

Bone fragments were mechanically ground in fine powder with a pestle in an agate mortar. 5 mg of bone powder was transferred to a 2 mL Eppendorf® tube (Eppendorf; Westbury, NY, USA) and 1 mL of demineralizing solution (5% trifluoroacetic acid (TFA) in water) was added. The solution was stirred during 24 h at 4 °C and then centrifuged for 10 min at 10,000 g at room temperature. The bone powder and the demineralizing solution were carefully separated. 1 mL of pure water was added to the centrifuged bone powder and the solution was stirred for 10 min at 4 °C. The solution was centrifuged for 5 min at 10,000 g, then the supernatant was removed. This step was repeated a second time. The delipidation was performed by adding 100 μL of water to the bone powder, then 900 μl of a chloroform / methanol solution (2/1 v/v). The sample was stirred for 1 h at 4 °C and then centrifuged for 10 min at 10,000 g. The lower phase that contains the lipids was removed. The bone powder was washed again twice with water. Residual water in bone powder was then evaporated at room temperature with a SpeedVac™ Concentrator (Eppendorf^TM^ Concentrator Plus, Eppendorf, Hamburg, Germany). The recovered demineralizing solution was evaporated at room temperature with a SpeedVac™. Then 200 μL of lysis buffer (8 M urea, 4 % SDS, 0.2% DCA, 50 mM DTT, 100 mM ammonium bicarbonate pH 8.8) were added to the bone powder and on the evaporated demineralisation solution. The suspensions were incubated overnight with shaking at 4 °C, before the eFASP digestion step.

### 2.4 eFASP digestion

Before use, 0.5 mL Amicon® ultra centrifugal filters with a cut-off of 10 kDa (EMD Millipore, Darmstadt, Germany) were freshly incubated overnight with the passivation solution containing 5% (v/v) Tween®-20. Amicon® ultra centrifugal filters filled with 200 μL water were poured in a water bath for 20 min each four times. The bone powder suspension and the demineralized fraction were transferred to separate Amicon® filters, 100 μL of exchange buffer was added (8 M urea, 0.2% DCA (deoxycholic acid) w/w, 100 mM ammonium bicarbonate pH 8.8) and then Amicon® filters centrifuged for 30 min at 10,000 g. The filtrates were discarded. 200 μL microliters of exchange buffer were added again into the Amicon® filters which were centrifuged. This step was repeated twice. The proteins were alkylated during 1 h at room temperature in the dark using 100 μL of alkylation buffer (8 M urea, 50 mM iodoacetamide, and 100 mM ammonium bicarbonate, pH 8.8). The Amicon® filters were centrifuged for 30 min at 10,000 g and the filtrate was discarded. After the alkylation step, 200 μl of exchange buffer was added to the Amicon® filter which was centrifuged for 30 min at 10,000 g and the filtrate was discarded. Two hundred microliters of digestion buffer (0.2% DCA (deoxycholic acid) w/w, 50 mM ammonium bicarbonate pH 8.8) were added to the Amicon® filter and then centrifuged. This step was repeated twice, discarding the filtrate at each step. The Amicon® filters were transferred to new 2 mL tubes. 100 μL microliters of digestion buffer and 40 μl of trypsin / LysC (Promega, Madison, USA) were added and incubated with shaking in a heating block tube (MHR23, Hettich, Netherlands) overnight at 37 °C. After this step, the peptides present in the Amicon® filter were recovered in the lower tube by centrifugation for 15 min at 10,000 g. In order to obtain a maximum of peptides, the Amicon® filters were washed twice with 50 μl of ammonium bicarbonate solution (50 mM pH 8.8). The filtrates containing all the peptides were transferred to 1.5 mL Eppendorf® tubes. 200 μl microliters of ethyl acetate and 2.5 μL of TFA were added inducing the appearance of a white precipitate. At once, 800 μL of ethyl acetate were added again and the Eppendorf® tubes were centrifuged for 10 min at 10,000 g. The organic phase was eliminated. This step was repeated twice. The Eppendorf® tube were placed for 5 min at 60 °C in a heating block tubes (SBH130, Stuart, Staffordshire, UK) to remove the remaining ethyl acetate. The samples were evaporated to dryness at room temperature with a SpeedVac™ Concentrator. The wall of the Eppendorf® tubes were rinsed with 100 μl of a methanol / water (50/50 v/v) mixture and again evaporated to dryness. For mass spectrometry analysis, the sample was dissolved again in 10 μl of water solution containing 0.1% of formic acid. The concentration was then estimated by measuring the OD at 215 nm using 1 μl of the solution using a droplet UV spectrometer (DS-11+, Denovix, Wilmington, USA). The samples were diluted at a concentration of 1 μg/μL before analysis.

### 2.5 Liquid chromatography-tandem mass spectrometry

LC-MS/MS analyses were performed on an Orbitrap Q Exactive plus mass spectrometer hyphenated to a U3000 RSLC Microfluidic HPLC System (ThermoFisher Scientific, Waltham, Massachusetts, USA). 1 μl of the peptide mixture at a concentration of 1 μg/μL was injected with solvent A (5% acetonitrile and 0.1% formic acid v/v) for 3 min at a flow rate of 10 μl.min^−1^ on an Acclaim PepMap100 C18 pre-column (5 μm, 300 μm i.d. × 5 mm) from ThermoFisher Scientific. The peptides were then separated on a C18 Acclaim PepMap100 C18 reversed phase column (3 μm, 75 mm i.d. × 500 mm), using a linear gradient (5-40%) of solution B (75% acetonitrile and 0.1% formic acid) at a rate of 250 nL.min^−1^ in 160 minutes and then 100% of solution B in 5 minutes. The column was washed for 5 minutes with buffer B and then re-equilibrated with buffer A. The column and the pre-column were placed in an oven at a temperature of 45 °C. The total duration of the analysis was 180 min. The LC runs were acquired in positive ion mode with MS scans from *m/z* 350 to 1,500 in the Orbitrap mass analyser at 70,000 resolution at *m/z* 400. The automatic gain control was set at 1e106. Sequentially MS/MS scans were acquired in the high-energy collision dissociation cell for the 15 most-intense ions detected in the full MS survey scan. Automatic gain control was set at 5e105, and the normalized collision energy was set to 28 eV. Dynamic exclusion was set at 90 s and ions with 1 and more than 8 charges were excluded.

### 2.6 Data analysis

Proteomics data were processed with Mascot (version 2.5.1, Matrix Science, London, UK) against NCBI database (NCBiProt_20190815) restricted to Mammalian (5,242,539 sequences). Error tolerant searches with up to three missed cleavages, 10 ppm mass error for MS and 15 mmu for MS/MS. Cysteine carbamidomethylation was set as fixed modification. Methionine oxidation and asparagine, glutamine deamidation were selected as variable modifications. The automatic error tolerant search which search the modified forms of already identified peptides was allowed. The second bioinformatics analysis was performed with PEAKS X (Bioinformatics software, Waterloo, Canada) against a home-made database containing 1,765 collagen sequences extracted from NCBi database (All_Collagen, downloaded 08-2019) restricted to Mammalian. Precursor’s mass tolerance was fixed to 10 ppm and fragment ion mass tolerance to 0.02 Da. Three missed cleavages were allowed. The same post-translational modifications (PTMs) above were allowed plus hydroxylation of amino acids (RYFPNKD) as variable modifications. Five variable PTMs were allowed per peptides. PEAKS PTM and SPIDER ran with the same parameters. Results were filtered using the following criteria: protein score −10logP ≥ 20, 1% peptide False Discovery Rate (FDR), PTM with Ascore = 20, mutation ion intensity = 5% and *Denovo* ALC ≥ 50%. Peptides with amino acids substitutions was filtered with minimal intensity set as 1E+7. Peptides identified on collagen I alpha 1 and I alpha 2 were aligned against the NCBI non redundant protein sequence (all non-redundant GenBank CDS translations+PDB+SwissProt+PIR+PRF excluding environmental samples from WGS projects containing 308,570,119 sequences) to find similarity with BLASTp (https://blast.ncbi.nlm.nih.gov/Blast.cgi?PAGE=Proteins). The scoring parameter alignment used BLOSUM62 matrix. The specific peptides were validated with a score of 100% of identity and full query coverage. Fragment ion spectra for specific peptides, deamidated and oxidative peptides and novel amino acid substitution biomarkers were manually examined for quality (Supplementary data 3, 5 and 6). The mass spectrometry proteomics data have been deposited on the ProteomeXchange Consortium (http://proteomecentral.proteomexchange.org) via the PRIDE partner repository (Vizcaíno, et al., 2012) with the data set identifier PXD021171.

## 3. Results

### 3.1 Bone samples

The bones and artefacts studied in this paper were recovered in Scladina Cave located 400 m southwest of the village of Sclayn (Andenne, Namur, Belgium), close to the Meuse River (Fig. 1). The site has been under scientific excavation since 1978. The cave infilling indicates a sequence covering at least the end of the Middle Pleistocene up to the Holocene (Pirson, et al., 2008, Pirson, et al., 2014). The cave stratigraphy indicates a sequence of the Upper Pleistocene (Pirson, et al., 2008). Lithic artefacts and faunal materials are found through the entire sedimentary sequence (Abrams, et al., 2010, Bonjean, et al., 2009). The fauna is composed of herbivorous species including horse (*Equus ferus*), red deer (*Cervus elaphus*), fallow deer (*Dama dama*), chamois (*Rupicapra rupicapra*), aurochs or bison (*Bos primigenius* or *Bison priscus*), woolly rhinoceros (*Coelodonta antiquitatis*). Omnivorous species including cave bear (*Ursus spelaeus*), brown bear (*Ursus arctos*), red fox (*Vulpes vulpes*) and arctic fox (*Alopex lagopus*), as well as carnivorous species including cave lion (*Panthera spelaea*), panther (*Panthera pardus*), cave hyaena (*Crocuta crocuta spelaea*) and wolf (*Canis lupus*). In the cave modified bones with cutting marks, filleting marks or breakage damages were also recovered (Otte, et al., 2000).

**Fig. 1.**
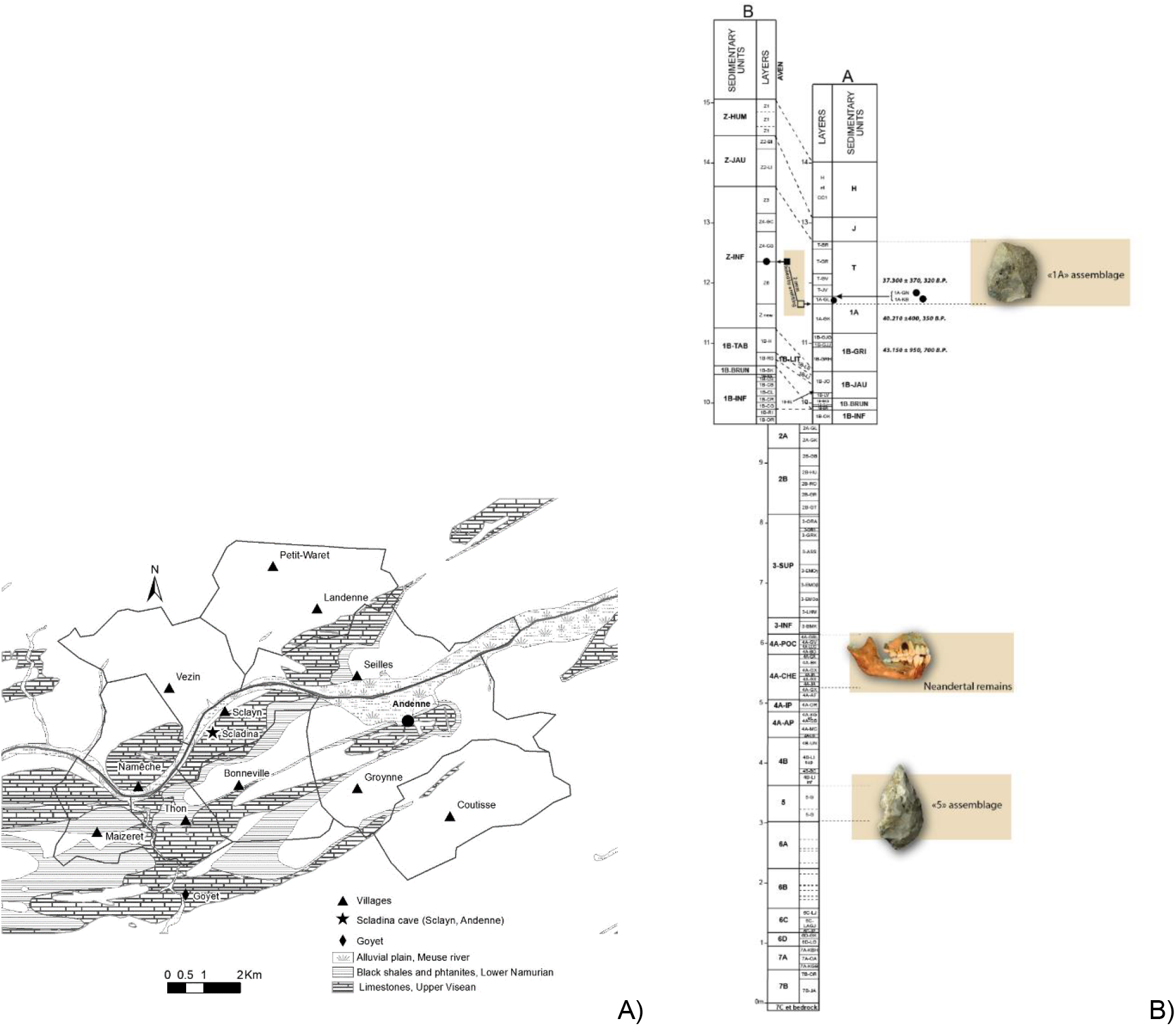
A) Location of Scladina Cave. Grey areas: distribution of the Palaeozoic carbonated rocks in Belgium. B) Stratigraphic profile of Scladina Cave composed of the entrance-sequence (lower and upper portions of A) and the aven sequence (upper portion of B; adapted from Bonjean et al 2015) (Bonjean, et al., 2015).

### 3.2 Proteomics methodology and identified proteins

We applied LC-MS/MS shotgun proteomics to ten selected 130,000 year old bones (Supplementary data 1). The method was applied to 5 mg starting material resulting in two analyzed fractions, the bone powder fraction and the demineralization residual fraction. Both fractions were digested with the eFASP method to increase the sensitivity, recovery and coverage of the identified proteins, and decrease the sensitivity to chemical contaminations (Erde, et al., 2014). As the trypsin enzyme is able to digest directly the proteins contained in bone powder (Schroeter, et al., 2019) no protein extraction was carried out on both fractions and trypsin was reacted directly on the bone powder and demineralization residual fractions. For bone powder fractions the mean Mascot score for identified peptides for the best hit score is 639 and 630 for demineralization residual fractions. The coverage sequence is roughly similar for both fractions 58 and 57 for the bone powder and demineralization residual fractions (Table 2). The Mascot score is very high for all identified proteins which tends to show a very good conservation of proteins inside the bones. For bone powder fractions the mean Mascot protein best hit score is 1618 and 1403 for demineralization residual fractions. The difference between the Mascot scores for both fractions is significantly different showing that more proteins are present in the bone powder fraction as expected as the demineralization step decalcifies the bone sample and remove the contaminants as keratins (Schroeter, et al., 2016). In our work, we used an aqueous 5% TFA demineralizing solution. TFA is a solvent commonly used in reverse-phase HPLC protein purification because of its effectiveness in solubilizing hydrophobic peptides. Using TFA, proteins or peptides are less modified as the TFA pKa is lower than HCl pKa which is the acid commonly used in paleoproteomics for demineralization (0.3 and −6.3 respectively). For all samples and fractions (i.e. bone powder and demineralization residual), the majority of the identified peptides matches mammalian collagen I alpha 1 (COL1A1) and collagen I alpha 2 (COL1A2) (Table 2). Considering COL1A1 and COL1A2 proteins, the peptides are distributed all along the sequences. The sequence coverage average for the best hit is 57% excluding the signal peptide and *N*, *C*-terminus propeptides. The other identified proteins with a low number of peptides are type II cytoskeletal 1 keratin, hornerin (less than 20 peptides for both proteins) which are contaminants of archaeological excavations (Hendy, et al., 2018). Others collagens are identified in both fractions for all samples as collagen II alpha 1 (COL2A1), collagen III alpha 1 (COL3A1), collagen V alpha 2 (COL5A2) and fibrillary collagen NC1 domain (Supplementary data 1).

**Table 2.**
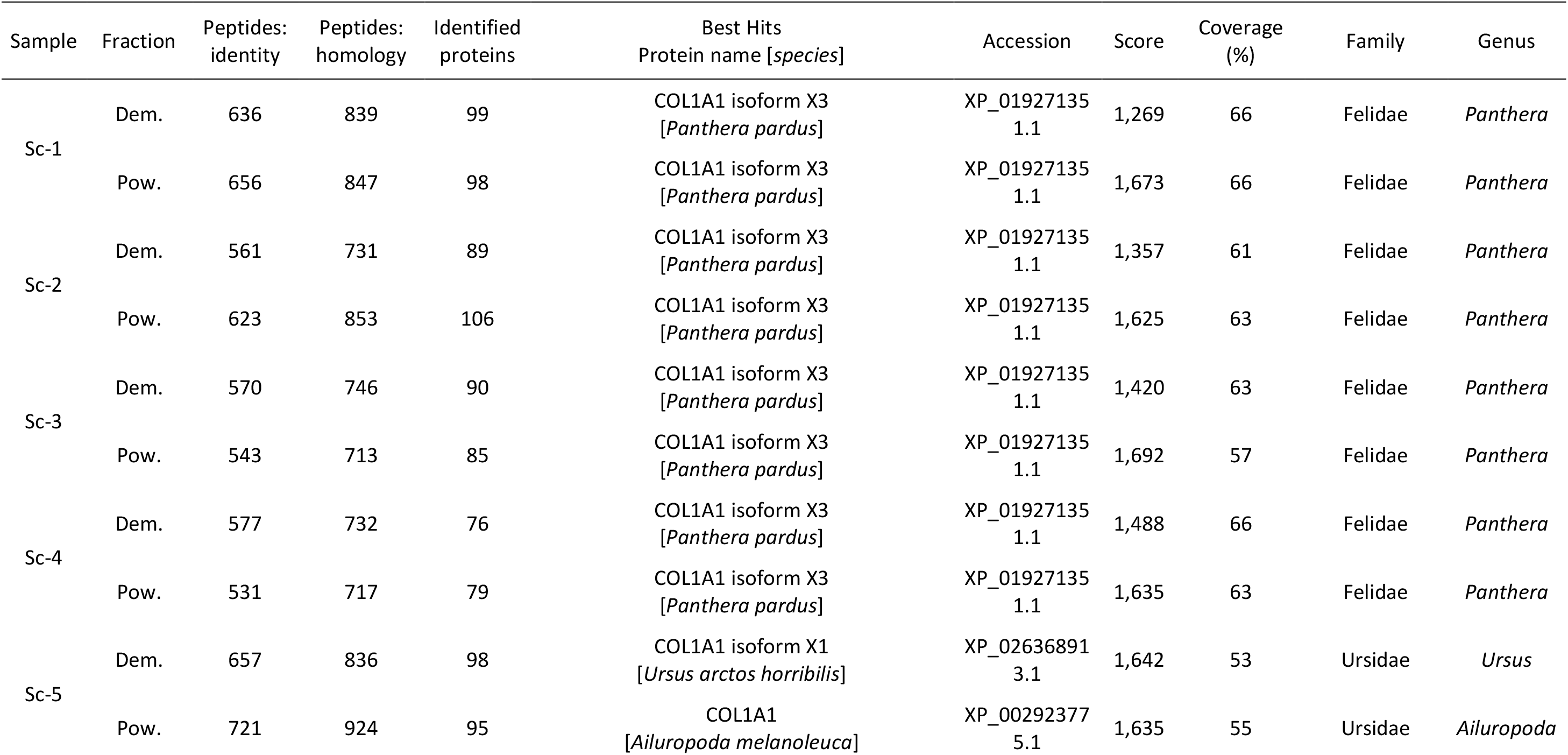
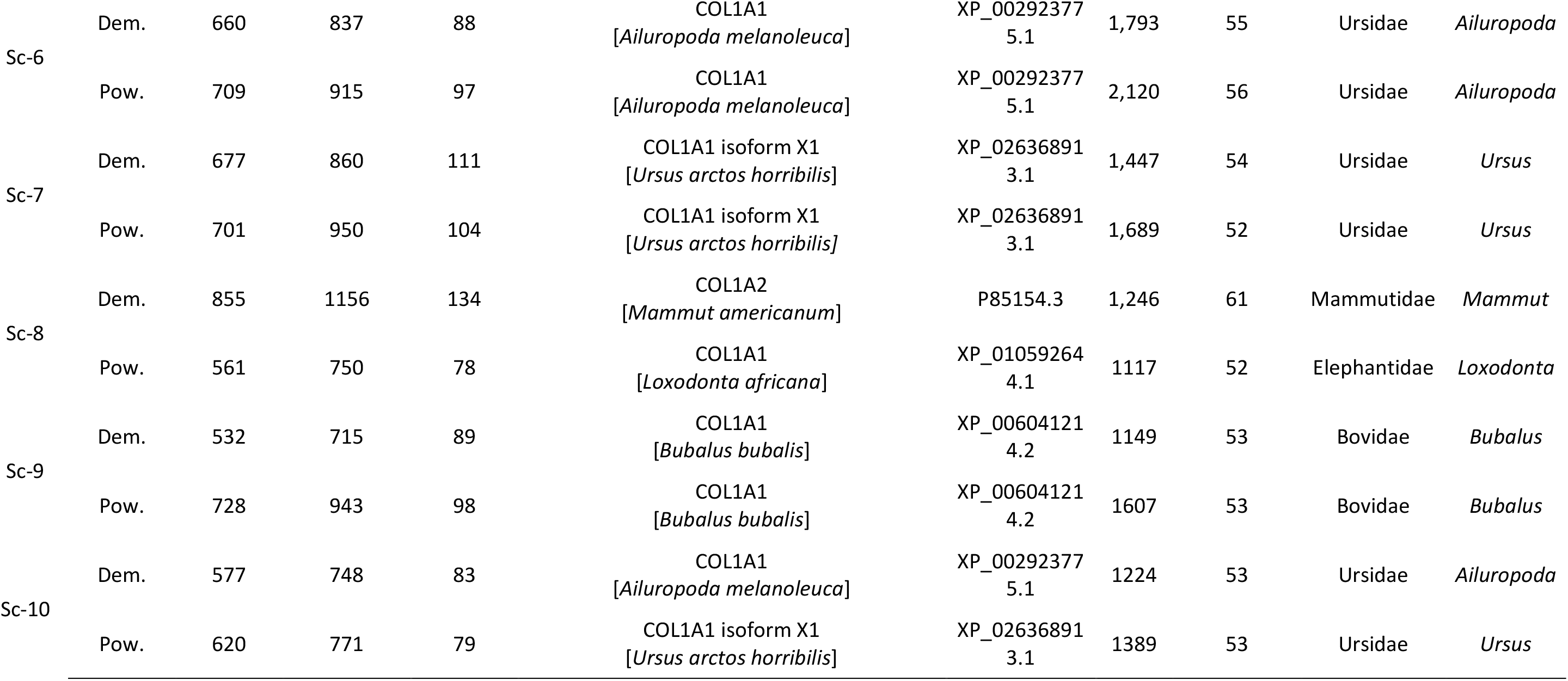
Information related to the quality of MS analysis for demineralization residual fraction (Dem.) and bone powder fraction (Pow.), including number of peptide matches above identity and homology thresholds, number of identified proteins, best hit and its NCBI accession number, Mascot protein score, best hit protein percentage of coverage. Family and genus taxa identified in archaeological samples (see sections 3.3 and 3.6). All identified proteins are listed in Supplementary data 2.

| Sample | Fraction | Peptides:<br>identity | Peptides:<br>homology | Identified<br>proteins | Best Hits<br>Protein name [*species*] | Accession | Score | Coverage<br>(%) | Family | Genus |
| --- | --- | --- | --- | --- | --- | --- | --- | --- | --- | --- |
| Sc-1 | Dem. | 636 | 839 | 99 | COL1A1 isoform X3<br>[*Panthera pardus*] | XP_01927135<br>1.1 | 1,269 | 66 | Felidae | *Panthera* |
|  | Pow. | 656 | 847 | 98 | COL1A1 isoform X3<br>[*Panthera pardus*] | XP_01927135<br>1.1 | 1,673 | 66 | Felidae | *Panthera* |
| Sc-2 | Dem. | 561 | 731 | 89 | COL1A1 isoform X3<br>[*Panthera pardus*] | XP_01927135<br>1.1 | 1,357 | 61 | Felidae | *Panthera* |
|  | Pow. | 623 | 853 | 106 | COL1A1 isoform X3<br>[*Panthera pardus*] | XP_01927135<br>1.1 | 1,625 | 63 | Felidae | *Panthera* |
| Sc-3 | Dem. | 570 | 746 | 90 | COL1A1 isoform X3<br>[*Panthera pardus*] | XP_01927135<br>1.1 | 1,420 | 63 | Felidae | *Panthera* |
|  | Pow. | 543 | 713 | 85 | COL1A1 isoform X3<br>[*Panthera pardus*] | XP_01927135<br>1.1 | 1,692 | 57 | Felidae | *Panthera* |
| Sc-4 | Dem. | 577 | 732 | 76 | COL1A1 isoform X3<br>[*Panthera pardus*] | XP_01927135<br>1.1 | 1,488 | 66 | Felidae | *Panthera* |
|  | Pow. | 531 | 717 | 79 | COL1A1 isoform X3<br>[*Panthera pardus*] | XP_01927135<br>1.1 | 1,635 | 63 | Felidae | *Panthera* |
| Sc-5 | Dem. | 657 | 836 | 98 | COL1A1 isoform X1<br>[*Ursus arctos horribilis*] | XP_02636891<br>3.1 | 1,642 | 53 | Ursidae | *Ursus* |
|  | Pow. | 721 | 924 | 95 | COL1A1<br>[*Ailuropoda melanoleuca*] | XP_00292377<br>5.1 | 1,635 | 55 | Ursidae | *Ailuropoda* |
| Sc-6 | Dem. | 660 | 837 | 88 | COL1A1<br>[*Ailuropoda melanoleuca*] | XP_00292377<br>5.1 | 1,793 | 55 | Ursidae | *Ailuropoda* |
|  | Pow. | 709 | 915 | 97 | COL1A1<br>[*Ailuropoda melanoleuca*] | XP_00292377<br>5.1 | 2,120 | 56 | Ursidae | *Ailuropoda* |
| Sc-7 | Dem. | 677 | 860 | 111 | COL1A1 isoform X1<br>[*Ursus arctos horribilis*] | XP_02636891<br>3.1 | 1,447 | 54 | Ursidae | *Ursus* |
|  | Pow. | 701 | 950 | 104 | COL1A1 isoform X1<br>[*Ursus arctos horribilis*] | XP_02636891<br>3.1 | 1,689 | 52 | Ursidae | *Ursus* |
| Sc-8 | Dem. | 855 | 1156 | 134 | COL1A2<br>[*Mammut americanum*] | P85154.3 | 1,246 | 61 | Mammutidae | *Mammut* |
|  | Pow. | 561 | 750 | 78 | COL1A1<br>[*Loxodonta africana*] | XP_01059264<br>4.1 | 1117 | 52 | Elephantidae | *Loxodonta* |
| Sc-9 | Dem. | 532 | 715 | 89 | COL1A1<br>[*Bubalus bubalis*] | XP_00604121<br>4.2 | 1149 | 53 | Bovidae | *Bubalus* |
|  | Pow. | 728 | 943 | 98 | COL1A1<br>[*Bubalus bubalis*] | XP_00604121<br>4.2 | 1607 | 53 | Bovidae | *Bubalus* |
| Sc-10 | Dem. | 577 | 748 | 83 | COL1A1<br>[*Ailuropoda melanoleuca*] | XP_00292377<br>5.1 | 1224 | 53 | Ursidae | *Ailuropoda* |
|  | Pow. | 620 | 771 | 79 | COL1A1 isoform X1<br>[*Ursus arctos horribilis*] | XP_02636891<br>3.1 | 1389 | 53 | Ursidae | *Ursus* |

### 3.3 Taxa classification

All analyses were carried out blind, especially for taxa classification. The identification of taxa discriminant peptides of COL1A1 and COL1A2 was performed using BLASTp analysis against the non-redundant NCBI database. In particular, we used the peptides from the taxa identified by LC-MS/MS analyses. We used for (i) *Panthera* taxa, peptides identified from *Panthera tigris altaica, Panthera pardus, Felis silvestris catus, Acinonyx jubatus, Puma concolor* and *Lynx pardinus*; (ii) for *Ursus* taxa, peptides identified from *Ursus maritimus, Ursus arctos horribilis, Ailuropoda melanoleuca*, (iii) for *Loxondota* taxa peptides from *Loxodonta Africana; Mammut americanum* and (iv) for the *Bos* taxa peptides from *Bos primigenius taurus, Bos grunniens mutus, Bos primigenius indicus, Bubalus Bubalis, Bison bison*.

Table 3 shows the unique peptides identified in the studied samples. Their corresponding MS/MS spectra are shown in Supplementary Data 3 (Fig. S3.1 – S3.19). All these unique peptides were identified in both fractions, i.e. demineralization residual and bone powder. Most of the identified peptides carry post-translational modifications like proline/methionine oxidation, glutamine/asparagine deamidation and carboxymethyl arginine. These modifications associated with protein biological ageing are found in analyzes of ancient bones (Cappellini, et al., 2011, Schroeter and Cleland, 2016, Wadsworth and Buckley, 2014).

**Table 3.**
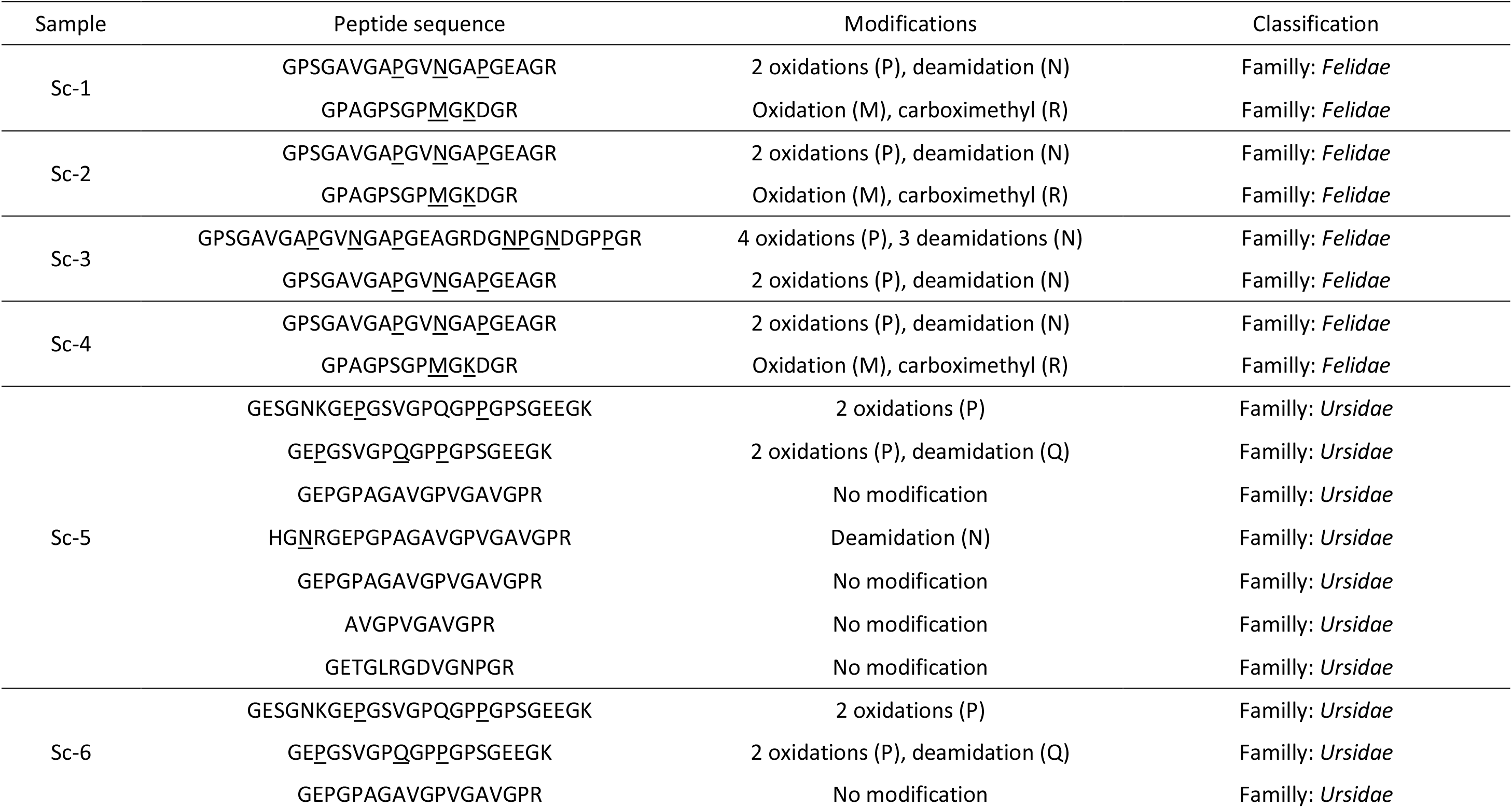
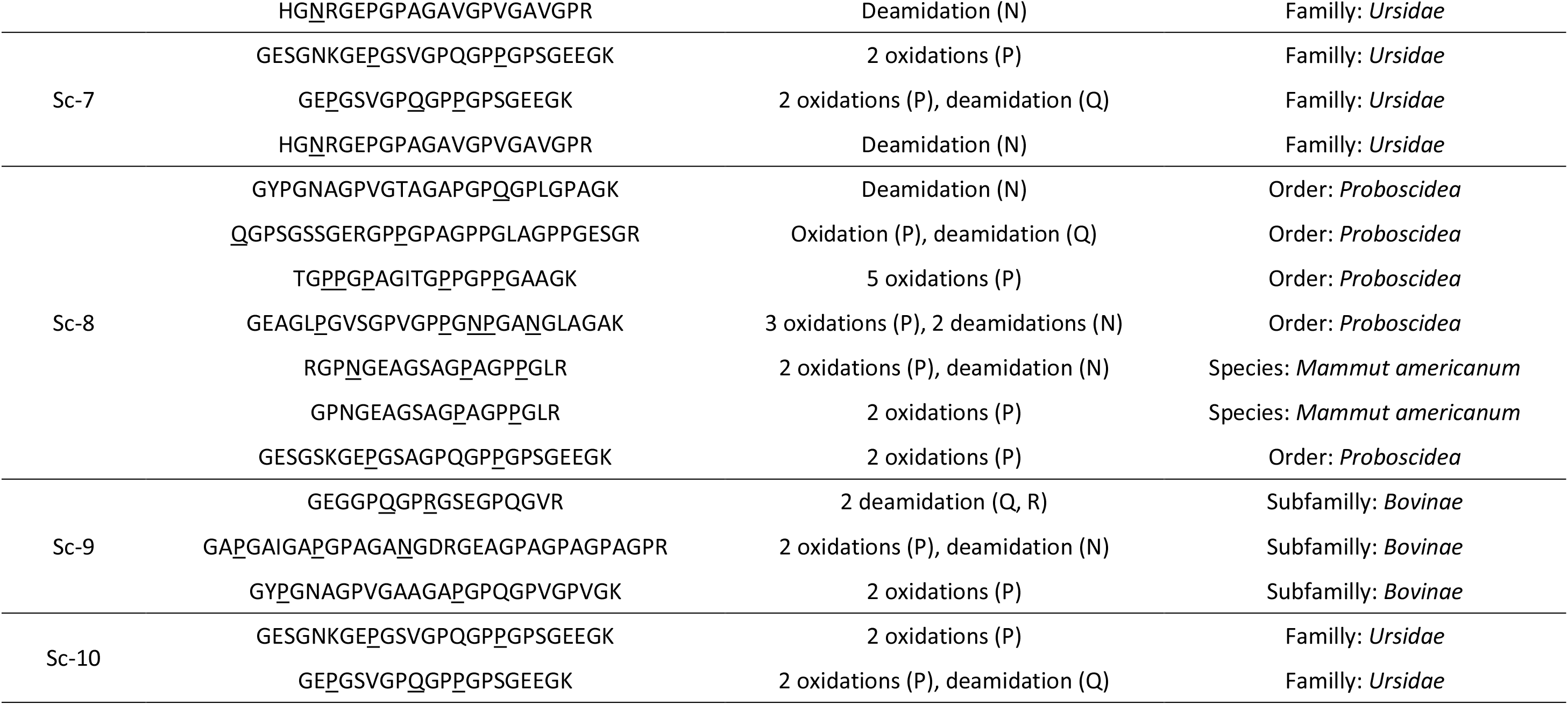
Unique peptides identified in the studied archaeological samples by LC-MS/MS. Underlined amino acids carry posttranslational modifications. The list of all the identified peptides and their modifications is given in Supplementary data 4.

Discriminant peptides matching the Ursidae family were identified in the samples Sc-5, Sc-6, Sc-7.

Considering the sample Sc-8, the unique peptides match the Proboscidea order and 2 unique peptides (RGPNGEAGSAGPAGPPGLR and GPNGEAGSAGPAGPPGLR) show specificity to *Mammut americanum* species. The Proboscidea order taxa classification corresponds to the morphological identification realized by palaeontologists.

Considering the Sc-9 and Sc-10 retouchers which were not identified by palaeontologist from the bone appearance, discriminant peptides for the Bovinae subfamily matching the *Bos primigenius taurus, Bos mutus, Bos indicus, Bubalus Bubalis, Bison bison* species as well as discriminant peptides for Ursidae family were identified respectively in samples Sc-9 and Sc-10. It can be remarked that 2 unique peptides from the sample Sc-10 were found in the samples Sc-5, Sc-6 and Sc-7.

Considering the Sc-1, Sc-2, Sc-3 and Sc-4 samples, discriminant peptides for the Felidae family were identified (Table 3) however, the identified peptides match all the following species (except *Lynx pardinus*): *Panthera tigris altaica, Panthera pardus, Felis silvestris catus, Acinonyx jubatus* and *Puma concolor*. Samples Sc-1 to Sc-3 were identified by paleontologists as *Panthera spelaea* bones and Sc-4 as *Panthera pardus* bone. The phylogenetic data shows that their closest ancestors are *Panthera leo* (Barnett, et al., 2016). Proteomics results may be refined allowing to trace amino acid mutations by introducing phylogenetic information. The species of closest ancestors were used to screen amino acid mutations. In our case only modern *Panthera pardus* collagen sequences are present in the databases and not *Panthera leo* sequences. So for samples Sc-1 to 4, collagen amino acids variations has been be studied from modern *Panthera pardus* sequences. Table 4 shows the closest species in the databases for the 10 studied bones and the phylogenetic data for each species. For retouchers, the faunal data found on the site, information from paleontologists, the results of proteomic analyses (number of peptides, sequence coverages) and phylogenetic data allowed targeting the closest species.

**Table 4.** Identification of the studied bones taxa by paleontologists and proteomic analysis and their closest species in the public proteins databases.

| Sample | Identified taxa |  | Closest species in databases | Phylogenetic reference |
| --- | --- | --- | --- | --- |
|  | Anatomy<sup>a</sup> | Proteomics<sup>b</sup> |  |  |
| Sc-1 | *Panthera spelaea* | *Felidae* | *Panthera pardus* | (Barnett, et al., 2016) |
| Sc-2 | *Panthera spelaea* | *Felidae* | *Panthera pardus* |  |
| Sc-3 | *Panthera spelaea* | *Felidae* | *Panthera pardus* |  |
| Sc-4 | *Panthera pardus* | *Felidae* | *Panthera pardus* |  |
| Sc-5 | *Ursus spelaeus* | *Ursus* | *Ursus arctos horribilis* | (Veitschegger, et al., 2018) |
| Sc-6 | *Ursus thibetanus* | *Ursus* | *Ursus arctos horribilis* | (Wu, et al., 2015) |
| Sc-7 | *Ursus thibetanus* | *Ursus* | *Ursus arctos horribilis* |  |
| Sc-8 | *Mammuthus sp* | *Proboscidea* | *loxodonta africana* | (Rohland, et al., 2007) |
| Sc-9 | Unknown | *Bovinae* | *Bos primigenius*<br>*Taurus* | (Park, et al., 2015) |
| Sc-10 | Unknown | *Ursus* | *Ursus arctos horribilis* | (Veitschegger, et al., 2018) |

<sup>a</sup>Species. <sup>b</sup>Order, family, subfamily or genus.

### 3.4 Asparagine and glutamine deamidation

The analysis of posttranslational modifications such as asparagine and glutamine deamidation or proline oxidation informs on the state of preservation of bones. Ancient, or at least more damaged, samples have a higher level of deamidation (Hill, et al., 2015, Ramsøe, et al., 2020, Schroeter and Cleland, 2016, Van Doorn, et al., 2012, Welker, et al., 2015). To calculate the frequency of deamidation and oxidation in all samples, 2 distinct peptides (GQAGVMGFPGPK and GANGAPGIAGAPGFPGAR) common to all the identified sequences of COL1A1 were used (Supplementary data 5, table S1). The quantified peptide modifications are oxidation on methionine or proline (*m/z* increment: + 15.9949), deamidation on glutamine or asparagine (*m/z* increment: + 0.9840). The sequence, mass, retention and fragmentation spectra of the native and modified peptides are given in Supplementary data 5 (Fig. S5.1 – S5.6).

The frequency of modification is calculated according to the formula (1):

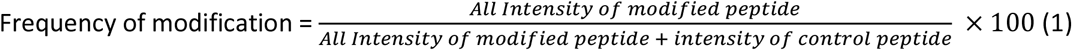

For COL1A1, in the bone powder fraction the average percentage of deamidation for all samples is 76.4% for glutamine and 85.5% for asparagine. These percentages are slightly higher in the demineralization residual fraction reaching 83.4% for glutamine and 91.4% for asparagine. The percentage of oxidation of prolines is stable at 99.1% for powder fractions and 99.4% for demineralization residual fraction (Supplementary data 5, Fig. S5.13). The same results are observed on COL1A2 (Supplementary data 5, Fig. S5.14). For COL1A1, in the bone powder fraction the average percentage of deamidation for all samples is 67.1% for glutamine and 80.7% for asparagine. These percentages are slightly higher in the demineralization residual fraction reaching 83% for glutamine and 80.9% for asparagine. The percentage of oxidation of prolines is stable at 99.6% for powder fractions and 99.7% for demineralization residual fraction (Supplementary data 5, Fig. S14). All samples exhibits elevated deamidation frequencies in the range expected from Middle Pleistocene bones (Lanigan, et al., 2020, Welker, et al., 2017). Pearson correlations show that the best correlation between bone powder fraction and the demineralization residual fraction is asparagine deamidation (*r* = 0.78 for COL1A1 and *r* = 0.62 for COL1A2) which proves that it is less impacted by the demineralization procedure.

### 3.6 Amino acids variation

Amino acid variants not referenced in the protein databases provide potential phylogenetic information. These variants may be found by error-tolerant search of MS/MS spectra, which is a computer intensive data processing. The very good conservation of proteins in bones from the Scladina Cave helped this study favorably. Supplementary data 6 (Fig. S6.1 – S6.56) shows the MS/MS spectra of peptides with amino acid substitutions for all samples and table 5 indicates the peptides with amino acids mutations. The analysis of variations of amino acids was carried out on the protein sequences of the species closest to the species identified by the paleontologists (Table 3).

**Table 5.**
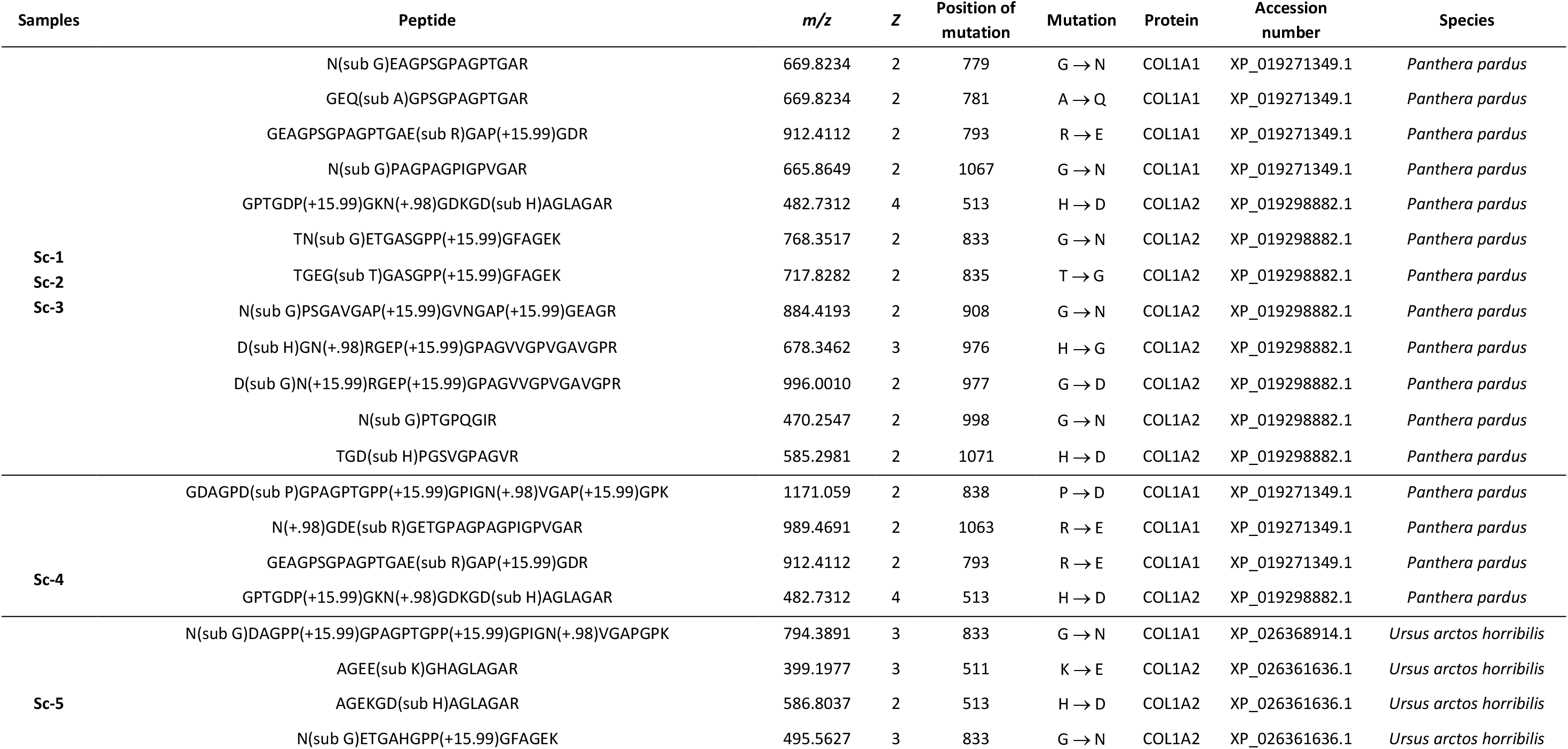
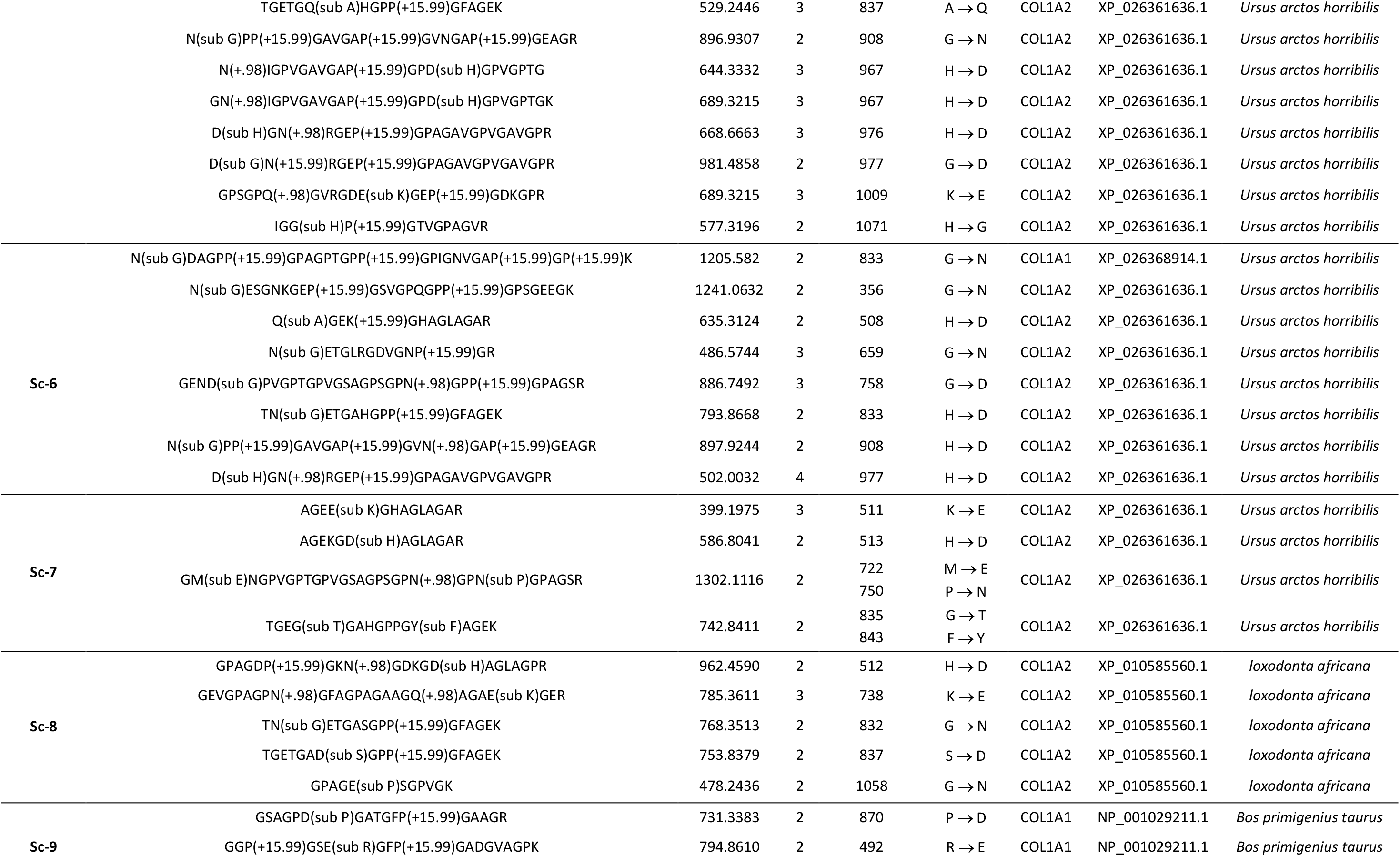
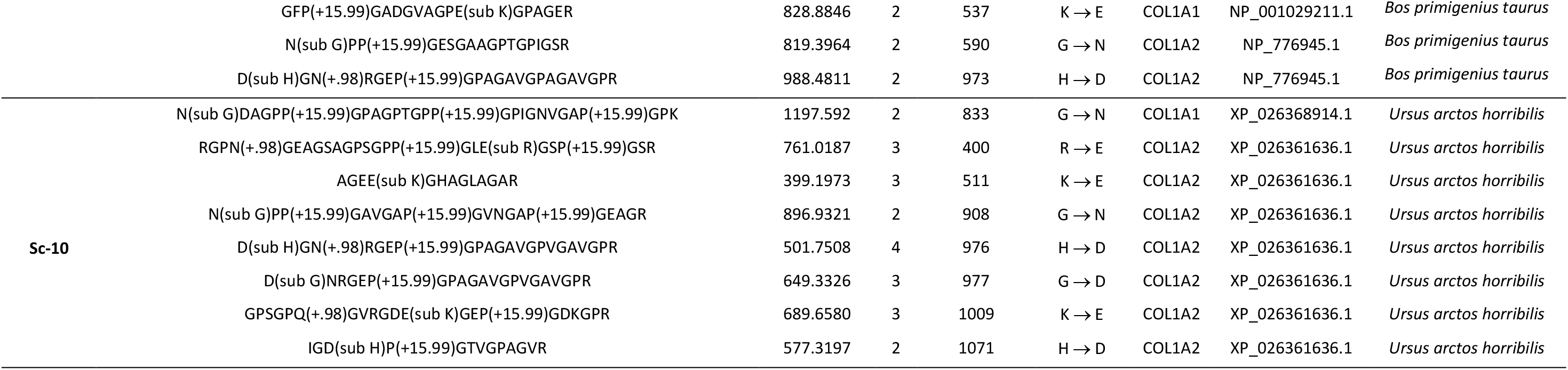
Peptides and amino acids variations identified in the studied archaeological samples. For each sample, the following information is described: peptides sequences, mass over charges (*m/z*), charge of peptides (*z*), the position and the type of mutation, the collagen chain, accession number and species. The peptides sequences are referencing the identified modifications (oxidation, deamidation, substitution). A *m/z* increase of +15.9949 corresponds to the oxidation of proline *m/z* increase + 0.9840 to deamidation of asparagine (N) or glutamine (Q). (sub *) denotes the substitution of amino acid where * is amino acid.

The samples Sc-1 to Sc-3 show twelve mutations on the sequence of collagens referencing to the *Panthera pardus* species. On the COL1A1 sequence, the substitution of arginine (R) by glutamic acid (E) is observed at position 793, the glycine (G) by asparagine (N) at position 779, the alanine (A) by glutamine (Q) at position 781 and the glycine (G) by asparagine (N) at position 1067. On the COL1A2 sequence, eight substitutions are observed at the positions 513, 833, 835, 908, 976, 977, 998 and 1071 (Table 5). On sample Sc-4, four mutations were detected; 3 on the COL1A1 sequence on position 793, 838 and 1063 and the last on the position 513 of the COL1A2 sequence (Table 5). The mutation on the COL1A1 sequence on position 793 and on the COL1A2 sequence on position 513 are also detected for samples Sc-1 to Sc-3 whereas the sample Sc-4 carries no other mutation. In brief samples Sc-1 to Sc-3 carries 10 identical amino acid variations in their protein sequences showing their belonging to the same species. These observations are in agreement with the osteomorphological analyses concluding to the identification of the *Panthera spelaea* species for the 3 samples, Sc-1 to Sc-3. The sample Sc-4 carries only 4 mutations in the collagen 1 alpha 1 and alpha 2 sequences compared to the modern *Panthera pardus* sequences. This observation is also in agreement with the identification made by the paleontologists.

The Sc-5 sample has been identified as a bone from the *Ursus spelaeus* species by paleontologists. The closest ancestor sequence in the databases is *Ursus arctos horribilis*. Proteomic analysis shows one amino acid substitution on the sequence of COL1A1 and eleven amino acid substitutions on the sequence of COL1A2 (Table 5). The Sc-6 and Sc-7 samples were identified as *Ursus thibetanus* species bones by paleontologists. Its closest ancestor is again *Ursus arctos horribilis*. One amino acid substitution on the sequence of COL1A1 and seven amino acid substitutions on the sequence of COL1A2 in Sc-6 sample. Four mutations in Sc-7 were observed on the sequence of COL1A2. There are no common mutations for samples Sc-6 and Sc-7. Only the mutations on the position 511 and 513 are common to the samples Sc-7 and Sc-5.

For the sample Sc-8, the closest ancestor is *Loxodonta Africana.* No substitution is observed on COL1A1 but 5 substitutions are observed on COL1A2 (Table 5).

The Sc-9 retoucher is identified as belonging to the *Bovidae* family. Considering the Scladina excavation site fauna, the closest animals are *Bos primigenius primigenius* (aurochs) and *Bison priscus* (bison). Collagen sequence of *Bos primigenius taurus* was identified with a greater number of peptides and a slightly higher sequence coverage. For COL1A1, *Bos primigenius taurus*: 408 peptides, 69% of coverage; *Bison* priscus: 149 peptides; 66% of coverage. For COL1A2, *Bos taurus*: 384 peptides, 71% of coverage; *Bison bison*: 243 peptides, 71% of coverage.

Three amino acid substitutions were also identified on the COL1A1 sequence and 2 substitutions on COL1A2 referring to *Bos Taurus* species. These results tend to show that the Sc-9 sample is a *Bos primigenius taurus* (aurochs) in agreement with Scladina Cave fauna.

Proteomics data indicates that the Sc-10 retoucher has been made from a bone of the *Ursidae* family. The closest species in protein database, referring to the Scladina Cave fauna is *Ursus arctos horribilis* as discussed above. Proteomics detected 1 mutations on COL1A1 sequence 7 mutations on COL1A2 sequence. Seven of these mutations are common with the Sc-5 sample (833, 511, 908, 976, 977, 1009 and 1071 positions). These results show a resemblance between the samples Sc-10 and Sc-5 potentially concluding that the Sc-10 sample may belong to the *Ursus spelaeus* species. The comparison between Sc-10 and Sc-6 and 7, show two mutation is in common with Sc-6 identified as at positions 833, 977 and one mutation is in common with Sc-7 identified as at positions 511. This results concluding that the samples Sc-6 and Sc-7 are not belonging to the same species than the sample Sc-10.

## 4. Discussion

### 4.1 Protein identification

Bone preservation and the presence of modern contaminants are two major difficulties of bone palaeoproteomics. In our study, the keratins were identified in minority in comparison to endogenous bone proteins like collagens. Human honerin, dermicin which are very often described as contaminants in bones proteomic analysis have been detected (Sawafuji, et al., 2017, Wadsworth and Buckley, 2014).

The two chains of the type I collagen, that represents 90% of the bone organic matter, were identified with up to 60% coverage for both chains. This result shows an exceptional bone preservation and a highly sensitive proteomic analysis based on an eFASP protocol (Erde, et al., 2014) without protein extraction after demineralization and 5 mg starting material. It can be pointed out that starting material ranging from 30 mg to several hundred mg are commonly referenced in palaeoproteomics studies (Buckley, et al., 2017a, Buckley, et al., 2017b, Buckley, et al., 2019, Cappellini, et al., 2011, Welker, et al., 2017)).

In this study, NCPs are not detected despite described in several ancient bones studies (Cappellini, et al., 2011, Sawafuji, et al., 2017, Wadsworth and Buckley, 2014) have been identified in our study. It is important to note that other collagens have been identified as the COL2A1 which is a fibrillar collagen found in cartilage, the COL3A1 which is found as a major structural component in hollow organs and the COL5A2 which is a minor component of connective tissue. Welker and colleagues showed the presence of others collagens as COL10A1 and COL27A1 in bones specimens from the Grotte du Renne (Welker, et al., 2016).

The LC-MS/MS of paleontological bone optimized eFASP digestion afford a high number of identified peptides, an important percentage of coverage and the identification of PTMs as very few spurious modifications are induced. The phylogenetic information on the species from the Scladina Cave afforded by paleontologists allow to identify the closest species in modern protein databases. By combining proteomic analysis, phylogenetic information and modern species protein data bases we identify the species of bones from the Scladina Cave which are in agreement with the paleontologists’ data. This methodology was applied to two retouchers Sc-9 and Sc-10 the species of which cannot be identified by their anatomy. Our study works on 5 mg of bone powder which is less than previously used quantities. This quantity is sufficient to identified proteins, genus, PTMs from Pleistocene bones, which indicates that our method is extremely sensitive.

### 4.2 PTMs

The preservation of bone proteins may be deduced from the percentage in weight of extracted collagen and the rate of deamidation which has been reported as a marker of degradation in ancient bone (Van Doorn, et al., 2012). Deamidation ranges from 0 to 20% for bones from 0 to 2000 years while older fossils show an extreme variance in their total levels of deamidation (Schroeter and Cleland, 2016). Deamidations of around 50% of glutamines and 60% of asparagine were found for 40 and 50 ka bones from the Kleine Fleldhofer Grotte (Germany) (Lanigan, et al., 2020). Considering middle Pleistocene bone specimens of the rhinoceros genus Stephanorhinus (337–300 ka), glutamine and asparagine deamidations reach nearly 100% (Welker, et al., 2017). In our study, the rate of deamidation ranges between 71 – 95% on the COL1A1 and 71 – 92% on the COL1A2 specific peptides identified in all Sc1-10 samples which is in agreement to the deamidation rates described in the literature (Welker, et al., 2017). The percentage of deamidation we observed in the demineralization residual fraction of collagen I alpha 1 is very close to the one observed in the bone power fraction.

Regarding proline oxidation, the collagen identified shows many peptides with hydroxylated prolines. The percentage of frequency is close to 99%, on the 60% of the collagen sequence which has been identified. In modern samples, the collagen prolines are oxidized at 20% (Zaitseva, et al., 2015). This modification is involved in stabilizing triple helix of collagen (Kotch, et al., 2008). The presence of peptides with hydroxyprolines can be explained by the age of the bones and the state of conservation. It is also interesting to notice that hydroxylysines were identified in the collagen sequences of Scladina Cave bones. Hydroxylysines are involved in crosslinks between the triple helix of collagen (Knott and Bailey, 1998). Hydroxylysines can be further modified by the sequential steps of *O-*linked glycosylation producing G-Hyl (galactosylhydroxylysine) and GG-Hyl (glucosylgalactosylhydroxylysine) (Scott, et al., 2012) and play multiple roles in normal mammalian physiology and pathology. Ryan C. Hill et al., have shown the presence of hydroxylysine glucosylgalactosylation in the extinct *Bison latifrons* (Hill, et al., 2015). Dihydroxyproline were identified in Scaldina cave samples. This modification has been reported in siliceous cell walls of diatoms (Ehrlich, et al., 2010, Nakajima and Volcani, 1969, Surmik, et al., 2016). These authors suggested that hydroxylated amino acids could play a role in silicification of diatom cell walls. The silicification is a fossilization process whereby the organism is penetrated by silica that form on the cells and cell structures. Bonjean and colleague showed the presence of silicified materials in the area of the Scladina Cave, which could explain the presence of dihydroxylation of prolines on the collagen sequences (Bonjean, et al., 2015).

### 4.3 Molecular sequence variation

Ancient proteins are commonly identified in references to the closest (most often modern) species present in the database as shown for Pleistocene samples of *Raphus cucullatus* from modern birds (Horn, et al., 2019), *Stephanorhinus* from *Rhinoceros* (Cappellini, et al., 2019, Welker, et al., 2017) or *kangaroos* from modern Australian vertebrate species (echidna, wombat, red kangaroo…) (Buckley, et al., 2017a).

This study shows a successful identification of bone taxa from Scladina Cave using proteomics and paleontological data analysis. Among the taxa identified by proteomics are the Felidae family, Elephantidae family, the Bovinae subfamily, the Ursus genus. Amino acid substitutions on the collagens were identified providing new information on extinct species sequences and helping in taxonomy-based clustering.

The samples Sc-1 to Sc-3 were identified as belonging to the Felidae family. No species information was extracted from our dataset. Osteomorphological studies identified the *Panthera spelaea* species however, no collagens are referenced in the database for this species (only mitochondrial sequences) nor for the closest species *Panthera leo* (Barnett, et al., 2016). The second closest species is *Panthera pardus* but again, no discriminant peptide was identified for this species nor the other referenced species (*Acinonyx jubatus*, *Felis silvestris catus*, *Lynx pardinus*, *Panthera pardus*, *Panthera tigris altaica*, *Puma concolor)*. Same amino acid substitutions of the COL1A1 and COL1A2 sequenced were found for the 3 samples (by referenced to the *Panthera pardus* modern sequence) indicating their belonging to the same species. A fourth sample, Sc-4, identified as belonging to the Felidae family showed different collagen sequence mutations compared to Sc-1 to Sc-3 samples indicating its belonging to another species. The Sc-4 bone has been identified as *Panthera pardus* by the paleontologists. These observations are in agreement with the phylogenetic studies carried out on mithochondrial DNA of *Panthera spelaea* (Barnett, et al., 2016, Joachim, et al., 2004, Ma and Wang, 2015).

The samples Sc-5, Sc-6, Sc-7 were identified as belonging to the Ursidae family by proteomics. Osteomorphological studies identified the Sc-5 sample as belonging to the *Ursus spelaeus* and the Sc-6 and Sc-7 samples to *Ursus thibetanus* species. The closest ancestor sequence in the databases is *Ursus arctos horribilis* for both species. Amino acids variations identified for the 3 samples could not help in taxonomy-based clustering. For the two bones Sc-6 and Sc-7, there is no mutations in common but the paleontologists have identified by osteomorphological studies as being of the same species. This can show variability between the collagen protein sequences of the same species. It would be interesting to have the genome sequencing of the species *Ursus thibetanus* species to confirm the Osteomorphological studies.

The sample Sc-8 showed specificity to the *Mammut americanum* species using 2 discriminant peptides on COL1A2 sequence and other unique peptides correspond to the Proboscidea order. The identification of the Proboscidea order taxa classification is in full agreement with the osteomorphological identification realized by the palaeontologists. The specific peptides, which showed specificity to the *Mammut americanum* species, are in contradiction with osteomorphological and faunal studies. We show in this study of importanceof correlating the proteomics data with the palaeontologists studies and to work on the protein sequences of the closest species.

The retouchers (Sc-9 and Sc-10) could not be identified by osteomorphologic studies. Proteomics identified the Sc-9 retoucher as belonging to the *Bovidae* family with a higher responses to *Bos primigenius primigenius* (aurochs) (number of peptides, slightly higher sequence coverage) in agreement with Scladina Cave fauna. The Sc-10 retoucher was identified to belong to the genus *Ursus*. Whereas Sc-10 shows different mutations than those observed for the Sc-6 and Sc-7 samples, similarities in amino acid mutations to the Sc-5 sample tend to conclude a to the Sc-10 belonging to the *Ursus spelaeus* species. Abrams *et al*. highlighted the presence of bear bone retouchers used by the local Neanderthal community present on the Scladina site (Abrams, et al., 2014) which were used tools for lithic activities. This taxa identification helps to confirm the exploitation of bear carcasses (brown and cave bears), recognised by the presence of butchery marks which remains a rare event during the Middle Palaeolithic. The use of bear bones by Neandertals is not very current and few sites shows remains of brown or cave bears concerned as retouchers, as in Northern France at Biache-saint-Vaast (Moigne, et al., 2016). So, this protocol of analysis, with a very low quantity of bone used, appear to be of great interest for Palaeolithic prehistorians.

Overall, the taxa identified by proteomics for the 10 studied samples correlate with the osteomorphological, palaeoenvironmental and palaeodietary data (Bocherens, et al., 1999, Bocherens, et al., 1997).

## 5. Conclusion

Our study has demonstrated the possibility to identify taxa of 130,000 years old bones using an optimized proteomics method starting from 5 mg of samples. A large number of peptides was identified along with high average sequence coverages up to of 60% for collagen 1 alpha 1. The deamidation frequency of about 80% and oxidation frequency of about 99% were measured. Despite the restricted databases for several species, several taxa were identified including the Felidae family, Elephantidae, Bovinae subfamily, the *Ursus* genus. Our study has shown not referenced amino acids substitutions specific to the studied species. Proteomics allowed to identify the retouchers’ originating families, bovidae and ursidae at Scladina Cave at the beginning of the Upper Pleistocene by Neandertals. Considering the amino acid variations of the collagen sequences and paleontological studies, one retoucher was identified as potentially belonging to the *Ursus spelaeus* species.

## Acknowledgments

The authors acknowledge the IBiSA network for financial support of the USR 3290 (MSAP) proteomics facility. The mass spectrometers were funded by the University of Lille, the CNRS, the Région Hauts-de-France and the European Regional Development Fund. The authors thank the CNRS Mission pour l’Interdisciplinarité for the funding of the project ProtéoMamQuat (2017-2018). The authors warmly thank the Scladina cave excavation team for granting them the bone samples of this study and for sharing their palaeontological data.

## 8. Credit roles

Fabrice Bray, Christian Rolando, Caroline Tokarski, Patrick Auguste: Conceptualization, Methodology. Dominique Bonjean, Grégory Abrams, Kévin Di Modica, Patrick Auguste: Resources. Fabrice Bray, Stéphanie Flament: Investigation. Fabrice Bray, Christian Rolando, Caroline Tokarski, Patrick Auguste: Writing – Original Draft. Fabrice Bray, Dominique Bonjean, Grégory Abrams, Kévin Di Modica, Christian Rolando, Caroline Tokarski, Patrick Auguste: Writing – Review & Editing.

## 9. Appendix A. Supplementary data

The mass spectrometry proteomics data have been deposited to the ProteomeXchange Consortium via the PRIDE partner repository with the data set identifier PXD021171.

The supplementary data 1 corresponds to pictures of ten bones from the archaeological excavation of Scladina Cave. The supplementary data 2 are excel files which containing the identification of proteins with Mascot engine against the Mammalian database. The supplementary data 2 contains one excel file for protein identified in demineralizing fraction and one other file for protein identified in powder fraction. The upplementary data 3 corresponds to the fragmentation spectra of unique peptides identified and presented in table 3. The supplementary data 4 corresponds to excel files containing the identified peptides for ten bones and both fractions from the archaeological excavation of Scladina Cave. The supplementary data 5 corresponds to the fragmentation spectra of peptides with oxidation on methionine, proline, asparagine and glutamine deamidation. Supplementary data 5 contains the figures of deamidation and oxidation frequencies for peptides form COL1A1 and COL1A2 for both fractions. Supplementary data 6 corresponds to the fragmentation spectra of peptides with amino acids variations identified in the studied archaeological samples.

